# Microfluidic device integrating a network of hyper-elastic valves for automated glucose stimulation and insulin secretion collection from a single pancreatic islet

**DOI:** 10.1101/2021.12.24.474087

**Authors:** Clément Quintard, Emily Tubbs, Jean-Luc Achard, Fabrice Navarro, Xavier Gidrol, Yves Fouillet

**Author notes:** Corresponding author: Clément Quintard; CEA, LETI, DTBS, 17 Av. des Martyrs 38000 Grenoble, France, Bat. 42 - P. A301-05.

## Abstract

Advances in microphysiological systems have prompted the need for robust and reliable cell culture devices. While microfluidic technology has made significant progress, devices often lack user-friendliness and are not designed to be industrialized on a large scale. Pancreatic islets are often being studied using microfluidic platforms in which the monitoring of fluxes is generally very limited, especially because the integration of valves to direct the flow is difficult to achieve. Considering these constraints, we present a thermoplastic manufactured microfluidic chip with an automated control of fluxes for the stimulation and secretion collection of pancreatic islet. The islet was directed toward precise locations through passive hydrodynamic trapping and both dynamic glucose stimulation and insulin harvesting were done automatically *via* a network of large deformation valves, directing the reagents and the pancreatic islet toward different pathways. This device we developed enables monitoring of insulin secretion from a single islet and can be adapted for the study of a wide variety of biological tissues and secretomes.

## 1. Introduction

There is growing interest in the development of microfluidic platforms that allow manipulation of various biological tissues and control of stimuli for the study of complex biological processes**(Rothbauer et al., 2018)**. Recently, several studies have shown the value of microfluidic devices for applications as diverse as drug-induced cardiotoxicity**(Jung Kim et al., 2011)**, anticancer drug testing**(Yang et al., 2018)** or pathogen detection**(Sayad et al., 2016)**. These studies are very promising and highlight the unique advantages of this technology such as its robustness, multiplexing capabilities, low manufacturing and maintenance costs, and ability to precisely control small volumes of liquid as well of the microenvironment around cells to better mimic relevant physiological conditions.

Notably, the application of microfluidics to pancreatic islets perifusion devices constitutes a promising approach for the reasons mentioned above**(Glieberman et al., 2019; Nourmohammadzadeh et al., 2016; Schulze et al., 2017; Shaikh Mohammed et al., 2009; Xing et al., 2016)**. Building *in vitro* analytical tools is crucial to model the complexity of physiology and pathophysiology of pancreatic islets for a better understanding of its basic biology as well as for the screening of new drugs. While most common perifusion systems require pooling of multiple islets to achieve quantifiable insulin concentrations, minimizing the number of islets required for experiments using microfluidic platforms is important given the scarcity of these biological tissues. Several groups have developed microfluidic devices that ensure single-islet sensitivity to tackle this issue**(Bandak et al., 2018; Dishinger et al., 2009; Godwin et al., 2011; Misun et al., 2020; Yi et al., 2015)**. Despite this, adoption of microfluidic platforms in industry and hospitals has been relatively slow due in part to complex technical settings (ease of use, scalability, etc.)**(Ewart and Roth, 2021; Rafael Castiello et al., 2016)**.

To improve automation and parallelization of microfluidic devices, many efforts were put on developing a microfluidic architecture with automatic hydrodynamic spheroid trapping, some of these involve parallel microchannels acting as a filter**(Munaz et al., 2016)**, 3D biological objects sedimentation into microwells**(Astolfi et al., 2016)**, or micro-rotational flows**(Ota et al., 2011)**. The hydrodynamic trapping method, initially proposed by Tan and Takeuchi**(Tan and Takeuchi, 2007)**, is of particular interest considering the serpentine architecture accompanied by narrowed chambers functioning as traps, and was used for pancreatic islets**(Nourmohammadzadeh et al., 2013; Silva et al., 2013)** as well as for cancer spheroids**(Ruppen et al., 2014)**. Remarkably, Zbinden et al.**(Zbinden et al., 2020)** integrated such a microfluidic platform with a Raman microspectroscope system to track the insulin secretion kinetics of pseudo-islets. However, these trapping devices are generally not coupled with fluid handling set-ups allowing complex biological assays on-chip. In particular, in this serpentine architecture with U-cup shaped constrictions, the flow is mostly redirected to the loop branch once the trap sites are occupied. It is thus crucial to reconfigure the design to force the flow through the trap sites where the islets are located in order to collect the secretions with high sensitivity. Valve-based microfluidics can be seen as the answer to overcome this challenge**(Oh and Ahn, 2006; Thorsen et al., 2002)**, but the small deformations that can be obtained with conventional PDMS, are better suited for single cell devices**(Zhang et al., 2020)**. When working with large 3D cell aggregates such as pancreatic islets (ranging from 50 μm to 500 μm), the elastic deformation properties of PDMS are no longer sufficient. However, reducing locally the dimensions of the fluidic microchannels to integrate valves is not an option, not only because of the fragility of pancreatic islets that cannot be deformed, but also because it would lead to excessive pressure losses, compromising the trap functioning. Recently, new types of materials qualified as hyper-elastic have been used for the fabrication of flexible robots**(Wehner et al., 2016)**. One of these materials is Ecoflex 00-50, which has a 1000% elongation rate, allowing the integration of valves inside the fluidic microchannels were biological tissues of several hundred micrometers can flow.

Here, we developed a microfluidic platform that is compatible with the size range of pancreatic islets and features a fluidic circuit board allowing complex biological assays. A single human pancreatic islets was passively directed along the fluidic path of least resistance to a constriction zone. The functionality of the pancreatic islet in the chip was tested by a glucose stimulation insulin secretion (GSIS) assay. Glucose stimulation and harvesting of the secreted insulin were achieved in an automated way through the activation of a valve network within the chip. Overall, the device presented here displays an active and programmable valve-based microfluidic circuit around a functional pancreatic islet, using a hyper-elastic material overcoming classical limitations.

## 2. Materials and methods

### 2.1 Human pancreatic islets

Two sources of pancreatic islets were used in this study.

3D pancreatic islet microtissues were purchased from InSphero (Schlieren, Switzerland). Upon the arrival of pancreatic islet microtissues in 96-well GravityTRAPTM plates, the microtissues were treated according to the manufacturer’s protocol and maintained in human islet maintenance medium (InSphero) before use (1 to 3 days). Age= 63, sex female (donor 1).

Human pancreatic islets were provided by the Diabetes Cell Therapy Laboratory, Institute of Regenerative Medicine and Biotherapies, St Eloi Hospital, Montpellier. The islets were isolated from the pancreas of brain-dead donors. The procedure was approved by the National Agency for Medicines and Health Products Safety (ANSM) in accordance with the French bioethics law (Act No.2004-800 of August 6, 2004, as amended by Act No.2011-814 of July 7, 2011). These procedures were carried out with approval by appropriate regulatory authorities. Islets were cultured in CMRL 1066 medium, 1g/L Glucose (Gibco, ref.041-96907A) supplemented with 10% FBS, 10% Glutamax 100X (Gibco, ref.35050.038), 10mM Hepes (Gibco, ref.15630-057), 100mg.UI.mL-1 Penicillin, 100 μg.mL-1 streptomycin (Gibco, ref.15140-122), in a 5% CO2 humidified atmosphere at 37°C. Viability of cells upon isolation= 80-90%, age= 60, sex male (donor 2). Islets with diameters greater than the constriction of 200 μm and smaller than the channel’s size and width of 400 μm were selected upfront for all experiments.

### 2.2 Glucose stimulated insulin secretion (GSIS) assays

#### Static control

Human pancreatic islets were individually placed in a 96 well plate with ultra-low attachment surface (Corning, NY, USA), and incubated at 37°C, 95% O2, 5% CO2 for 1 hour in 150 μL KREBS buffer 1% BSA, 2.8 mM glucose as pre-incubation. Thereafter, islets were incubated for 1 hour with 2.8 mM glucose solution (low glucose solution, LGS) and for 1 hour with 16.7 mM glucose solution (high glucose solution, HGS). After each incubation step, supernatants were collected and stored at −80□°C. Insulin concentration in each collected supernatant was measured using STELLUX Chemiluminescence Human Insulin ELISA (ALPCO).

#### GSIS on-chip assay

Pancreatic islets were stimulated following a classic GSIS assay protocol adapted to a dynamic on-chip configuration. The chosen microfluidic architecture takes advantage of the control capabilities of the valves to direct the different reagents. Briefly, we performed a multi-step protocol using a network of valves integrated into the fluidic channels. First, a single human pancreatic islet was loaded into the chip through the main serpentine pathway at the flow rate of Q = 100 μL/min (Fig. 1a, green path). The low (Fig. 1a, yellow path) and high (Fig. 1a, red path) glucose stimulation were then performed *via* different pathways induced by the actuation of the valves network to direct the flow appropriately. Finally, the islet was retrieved from the chip *via* the main serpentine pathway for further KCl stimulation. All these phases lasted one hour. All operations were done into a thermostatic chamber at 37°C. A low flow rate of Q=2.5 μl/min was set for all the fully automated secretion phases.

**Fig. 1.**
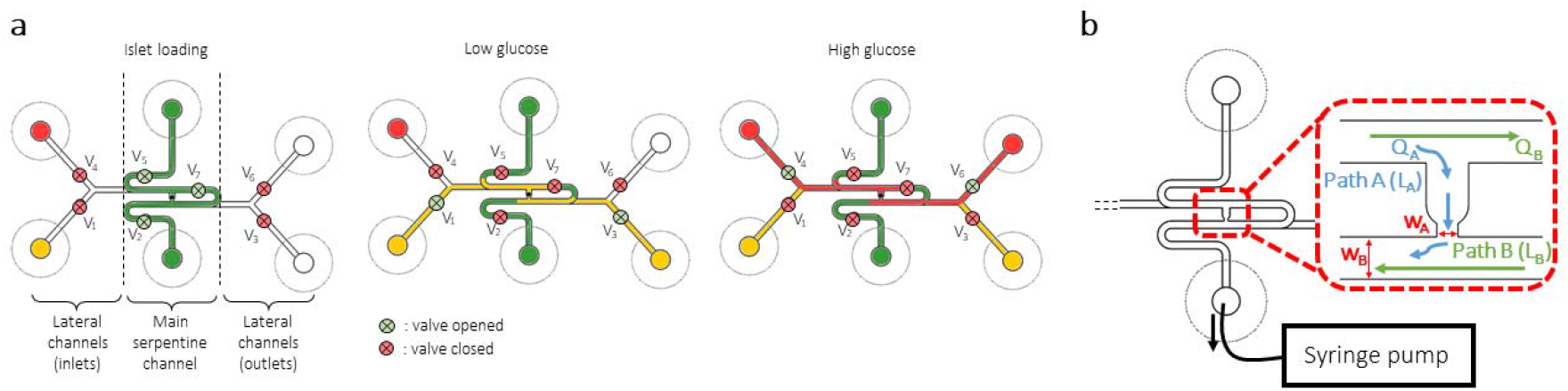
Glucose stimulated insulin secretion (GSIS) on-chip microfluidic architecture. a. Schematic diagram showing an overview of the GSIS assay on-chip protocol using a serpentine microfluidic architecture coupled with a network of pneumatic valves. b. Top view of the microfluidic main channel. A syringe pump was connected to the outlet of the channel to introduce the different reagents. Close-up on the U-cup shaped islet trap based on a hydrodynamic resistance difference between two possible flow paths (inset).

### 2.3 Trapping principle and islets loading

The microfluidic channel was composed of a serpentine-shaped channel with a narrow U-cup shaped region that functioned as a 3D-cell aggregate trap (Fig. 1b). When the trap was empty, the islet followed the path of least hydrodynamic resistance (path A), while an islet occupying the trap site allowed to bypass the U-cup shaped region and redirected the flow into the loop channel (path B). As a result, an islet in the flow was carried into the U-cup shaped constriction at a flow rate of Q = 100 μL/min where it was then protected from high shear stress (see Supplementary Note 1 for details). In order to precisely monitor the flow, the loading of different components was done through a syringe pump connected to the outlet of the channel in its withdrawal. Islet stimulation and secretions collection were achieved through the lateral channels housing at the entrance two reservoirs for low and high glucose solutions, and connected to the syringe-pump at the outlets.

### 2.4 Computational simulation of flow dynamics and shear stress

Fluid flow in the main serpentine channel was modelled with COMSOL Multiphysics 5.4 (COMSOL, Burlington, MA, USA) using the laminar flow module. The 3D geometry was imported from Solidworks, and a block was placed in the channel to mimic the valve forcing the flow through the U-cup trap. Islet porosity was omitted and the islet was represented by a solid sphere resting on the walls of the trap site. The properties of water were assigned to the fluid flowing at Q = 2.5 μL/min in the microchannel. A fine physics-controlled mesh was used for a stationary analysis. Shear stress on the islet surface was calculated for a 300 μm diameter islet as the product of the dynamic viscosity and the shear rate 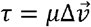.

### 2.5 Insulin quantification

Insulin concentrations were measured from culture supernatants collected during the GSIS assay, using the STELLUX^®^ Chemiluminescence Human Insulin ELISA (ALPCO) assay and the Tecan (Magellan) plate reader. Each sample was measured in duplicate. A stimulation index was obtained by calculating= [HGS]/[LGS].

### 2.6 Islet viability

The viability of the islets was determined after GSIS assay by a 15 min incubation in SYTO13 solution (final concentration= 2.5 μM) followed by a 5 min incubation in Propidium Iodide (PI) solution (final concentration= 10 μg/ml). Confocal z-stacks of the stained individual islets were taken and flattened in ImageJ to a 2D maximum intensity projection. The quantification of the nuclei of SYTO13-labelled live cells and the nuclei of PI-labelled dead cells allowed to assess the islet viability as follows: cell viability(%)=(number of green nuclei)/(number of green + red nuclei)* 100.

### 2.7 Imaging

Movies of fluorescent microbeads (Thermo Fisher Scientific Fluoro-Max Fluorescent Beads) infusion were taken at 15 Hz on an inverted IX50 microscope equipped with a CCD camera interfaced with Point Grey Software. Image processing was conducted using ImageJ (National Institute of Health, New York, NY, USA), codes are available on request.

### 2.8 Chip design and fabrication

In this study, we used hybrid microfluidic chips comprised of two or polymer layers and a stretchable membrane.

For glucose stimulated insulin secretion (GSIS) assays on-chip, we fabricated the microfluidic chip with a fluidic layer (F) and a pneumatic layer (P) enclosing the hyper-elastic membrane (Fig. 2a). Here, the pancreatic islet as well as the different reagents required for the GSIS assays flowed into the microchannels of the fluidic layer. Channels were micromachined in the pneumatic layer and positioned at different locations in the microfluidic circuit to create a valve network (Fig. 4a). These pneumatic channels were connected to a pressure source to push the membrane toward the fluidic channels. This enabled to open or close the fluidic channels at seven locations in the circuit functioning as valves. The valves were considered opened when no pressure was applied (P_A_), partly closed (or leaky) when an intermediate pressure P_I_ was applied and closed when a pressure above the critical pressure P_C_ required to deform the membrane by the height H of the fluidic channel was applied (Fig. 2b). Importantly, because the Ecoflex is a gas permeable material, the pneumatic channels were filled with liquid (PBS) prior the experiments to avoid the formation of bubbles in the fluidic channels when the valves were actuated. Finally, to ensure the tightness of the valves, the fluidic channels were machined with a round milling cutter to form hemicylindrical channels so that the membrane can perfectly match their shape. Pneumatic channels were machined to a depth and width of 200 μm. They were connected to the pressure controller via standard luer ports and the membrane was pierced with a needle at these specific areas to allow the air to flow toward the microchannels. Fluidic channels dimensions were 400 μm or 200 μm in height and 400 μm in width, and were connected to the syringe-pump via standard luers ports.

**Fig. 2.**
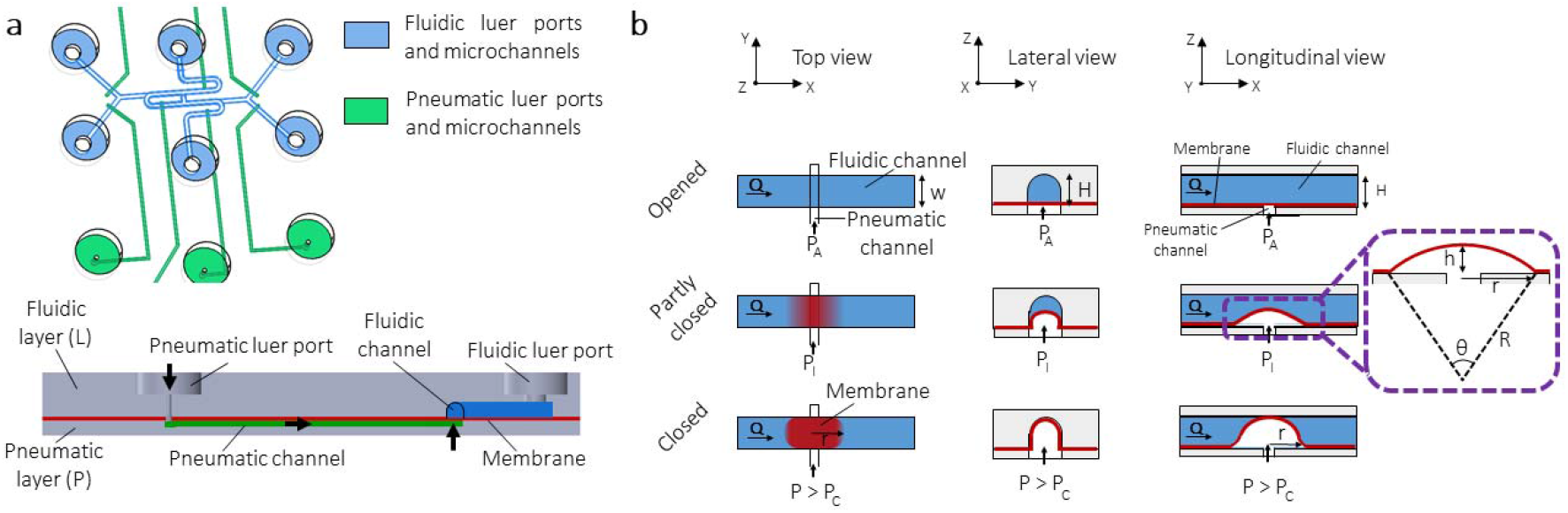
Overview of the operation principle of the hyper-elastic membrane. a. Solidworks 3D view (top) of the microfluidic chip, consisting of a hyper-elastic membrane enclosed between a fluidic layer and a pneumatic layer, and sectional view (bottom) showing the path taken to deform the membrane (black arrows). b. Schematic sectional views of the integrated hyperelastic valves on-chip in the three possible states (opened, partly closed, closed).

**Fig. 3.**
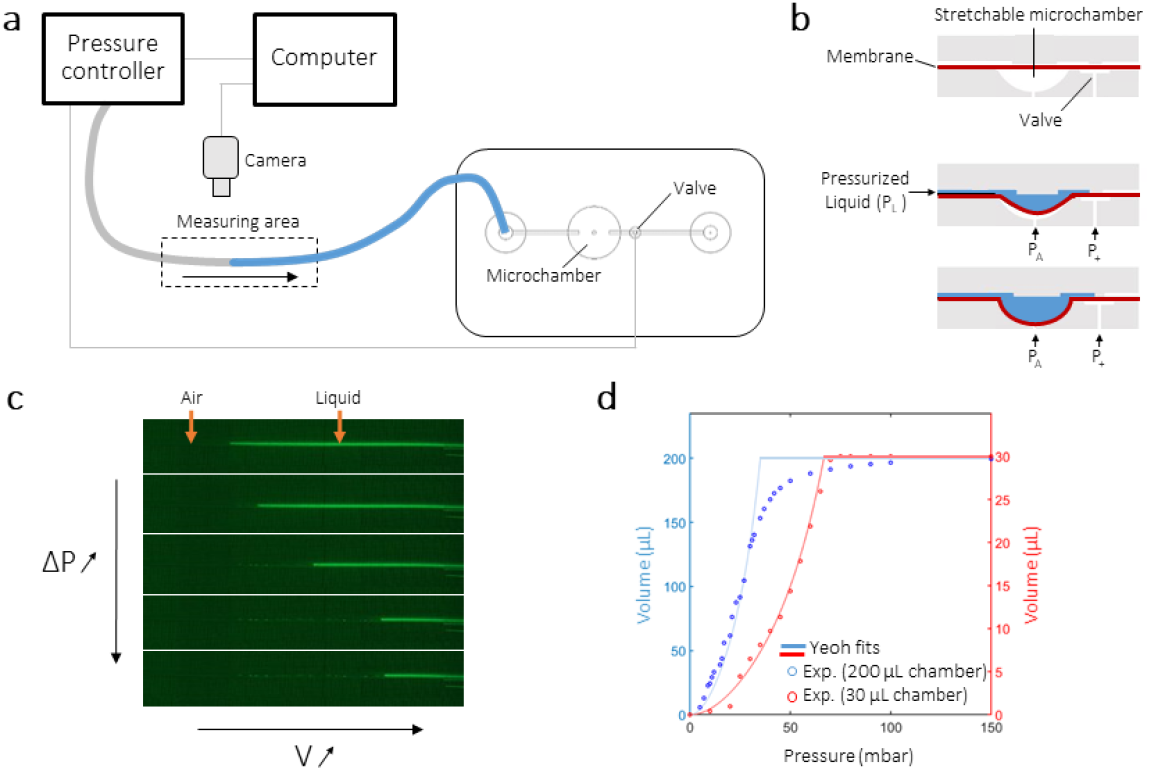
Determination of the volume of a stretchable microchamber. a. Schematic diagram of the experimental set-up. b. Schematic sectional view of the microchamber. The hyper-elastic membrane was stretched to its maximum capacity defined by the milling of the pneumatic layer. c. Measurement of the meniscus advancement as the pressure was increased using the pressure controller. d. Relationship between the volume of liquid injected in the microchamber and the pressure imposed by the pressure controller: comparison of the experimental results with the experimental results with the Yeoh law.

**Fig. 4.**
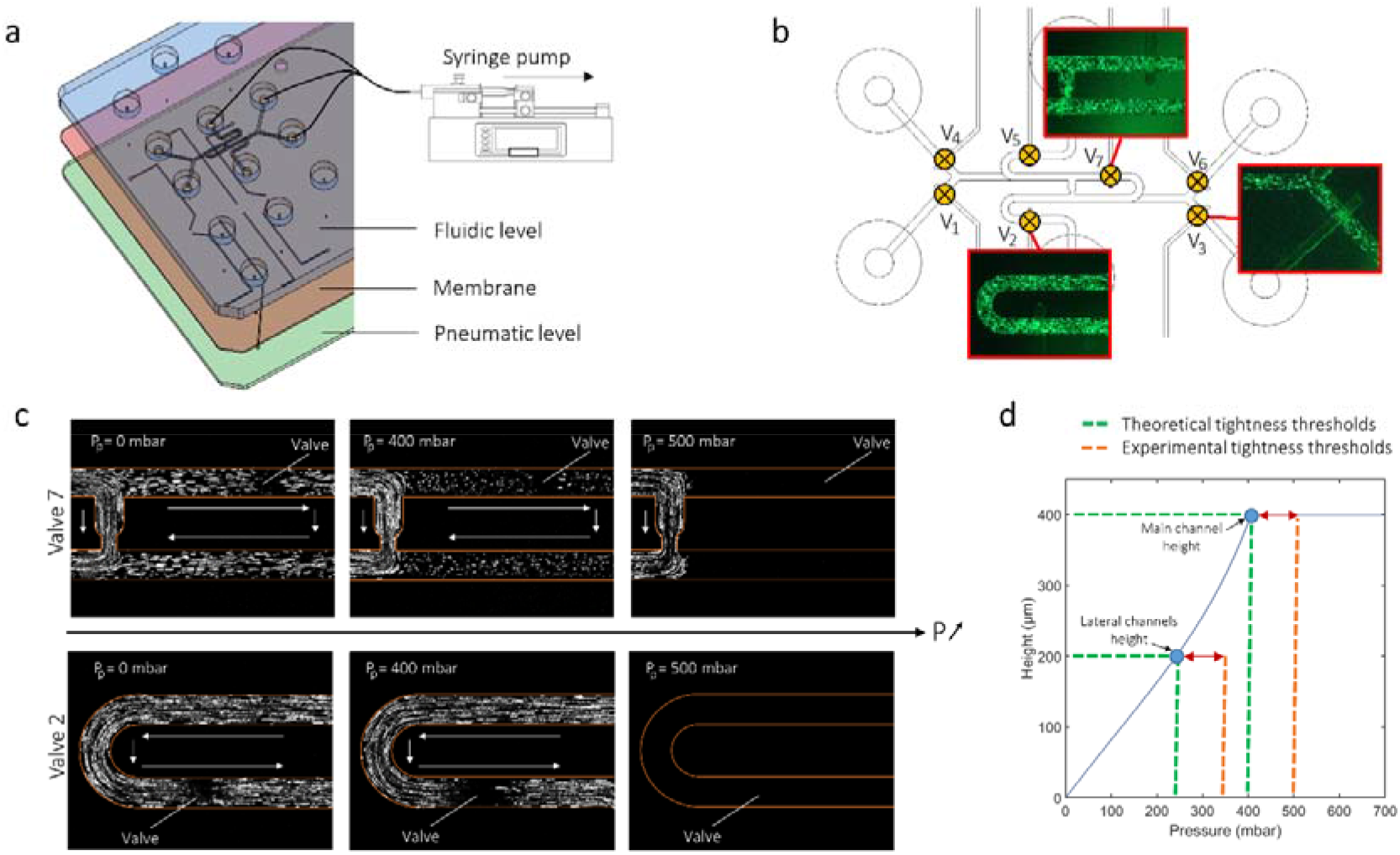
Valves validation in the GSIS assay chip. a. Solidworks 3D exploded view of the GSIS assay microfluidic chip. b. Top view of the fluidic microchannels and the locations of the valves. Fluorescent microbeads infusion were recorded around V_7_, V_2_ and V_3_ (red insets). c. Z-projection of maximum intensity over a 5 images stack, highlighting the tracks of the microbeads flowing through the microchannels. The stack was obtained after an image subtraction process. d. Comparison of the valves tightness pressure thresholds obtained experimentally and predicted by the Yeoh model.

To characterize the behavior of the hyper-elastic membrane when embedded in the microfluidic chip, experiments were conducted on a chip manufactured as previously described, comprising a set of elementary chambers whose volumes were controllable. Hemispherical chambers were micromachined on the P layer, as well as microchannels communicating with the valves and hemispherical chambers. This layer was placed facing the F layer covered by the membrane. Microchannels allowing communication with the stretchable chambers and the fluidic inlets and outlets, were micromachined on the F layer.

The microfluidic chips were designed using Solidworks software. Layers dimensions were credit card format (85.6 x 54 mm). With the aim of scalable manufacturing, microfluidic patterns were directly machined in a Cyclic Olefin Copolymer (COC) sheet (TOPAS, US) using a DATRON M7HP equipment (DATRON, GE), a material suitable for industrial production**(Nunes et al., 2010)**.

The elastic membrane was fabricated by spin coating Ecoflex 00-50 (Smooth on, US) on a silanized silicon wafer at 3000 rpm for 6.5 seconds, resulting in a ≈ 150 μm thick Ecoflex layer. This bi-component silicone material withstands larger deformations than PDMS, exhibiting a Young’s modulus around 200 kPa and a maximal stretching ratio before breaking of 980%. The membrane was plasma-bonded to the pneumatic and fluidic layers and fluidic layers following a protocol adapted from COC - PDMS bonding**(Cortese et al., 2011)**. Prior to bonding, the surfaces of COC polymers were cleaned with ethanol, rinsed in water and dried. The surfaces of the pneumatic and fluidic layers were activated by oxygen plasma and silanized by a 30 min immersion in a 2% 3-amino-propyl-triethoxy-silane (APTES) solution. The layers were then thoroughly rinsed in water and dried. The plasma activated Ecoflex was laminated on the pneumatic level, before being covered by the fluidic level. Finally, the assembly was press-sealed for 15 minutes (Supplementary Fig. 1).

### 2.9 Modelling of the hyper-elastic membrane deformation

The stress-deformation relation that was used here to describe the hyper-elastic membrane characteristics is the one derived from the Yeoh’s model**(Yeoh, 1993)**, a higher-order extension of the neo-Hookean law, commonly employed for incompressible, nonlinear elastic materials such as Ecoflex. To evaluate the validity of the model in our configuration and to determine the pressure levels required to activate the membrane until it obstructs the fluidic channels, experiments were conducted. In these studies, the mode of deformation being equibiaxial, the elongations λ_1_, λ_2_ and λ_3_ in the main directions of space have been simplified to λ_1_ = λ_2_ = λ and λ_3_ = λ^-2^. The expression of the mechanical stress was thus given by **(Rodríguez-Martínez et al., 2015)**:

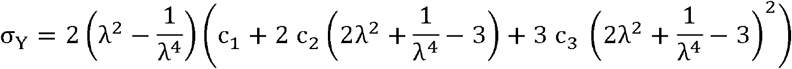

In these relations, the C_i_ are constants specific to each material. In the case of Ecoflex 00-50, C_1_, = 19000 Pa, C_2_ = 900 Pa and C_3_ = –4.75 Pa**(Xavier et al., 2021)**.

The results were then processed in a simplified way using by analogy the Laplace formula 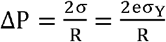 (Fig. 2b inset), where eσ_Y_ plays the role of the surface tension, e being the membrane thickness. The elongation 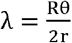 was expressed as a function of r and h using geometrical arguments**(Quintard et al., 2019)** (see Supplementary Note 2 for details). The satisfactory results have subsequently allowed to apply this model to the GSIS assay chip to estimate the pressures that would guarantee the tightness of the valves.

## 3. Results and discussion

### 3.1 Modelling the hyper-elastic membrane

To make the microfluidic valve network compatible with the size of the islets (ranging from 50 μm to 500 μm), another material than PDMS must be considered as it exhibits too low deformation rates. As a matter of fact, for the valves to be operative, the membrane must be deformed by the height of the channel, which itself must be larger than the pancreatic islets in order not to squeeze them. The high elasticity of the Ecoflex material allows very low actuation pressures, which is a real advantage in terms of simplification of the instrumentation and integration. However, the study of the deformation of such a material is classically carried out using blowing tests, but these do not correspond to our experimental on-chip configuration. Indeed, since Ecoflex is a flexible material, it is possible to have slippage when embedding it in the microfluidic chip. Moreover, before closing the microfluidic cards on the Ecoflex, the latter is pre-stressed for an optimal rolling process. It is therefore difficult to know if the results of a classical blowing test are still valid in such a device. Thus, a characterization of the membrane deformation directly on the microfluidic chip was necessary. To accurately study the behaviour of the membrane, we chose a hemispherical deformation model, which can easily be adapted to describe valve closure in the GSIS assay chip. The results of experiments measuring stress-strain relationships for soft materials have long shown that the linear theory of elasticity, in particular Hooke’s law used since the 17th century, was inappropriate for characterizing new materials with high elongation rates**(Mooney, 1940; Rivlin, 1948)**. A new theory, which has given rise to various models, is based on a strain energy density function**(Martins et al., 2006)**. We examined if the Yeoh law was able to describe our experimental configuration.

The experiment was conducted on the dedicated microfluidic chip consisting of a chamber fed by a capillary connected to a pressure controller (Fig. 3a). A valve was kept closed downstream during the filling of the chamber. The aim of the experiment was to measure the volume of liquid injected under the membrane for a given pressure. To do so, a camera filmed the advancement of the meniscus formed by the liquid in the capillary according to the pressure imposed by the pressure controller. A pressure was applied at the entrance of the tube by the pressure controller and increased by steps of 10 mbar while the advancement of the meniscus was measured at each step. The chamber was left at atmospheric pressure *P_A_* and the valve downstream of the chamber was kept closed, pressurized to *P*_+_ = 500 mbar (Fig. 3b). By measuring a relationship between the volume of liquid injected into the chamber and the pressure imposed by the pressure controller was thus found (Fig. 3c). Experiments were conducted on chambers of 30 μL volumes and 200 μL, for pressures ranging from 0 mbar to 70 mbar and 150 mbar respectively. Fig. 3d shows a strong agreement between theoretical and experimental curves, thus validating our model of deformation of the membrane in the microfluidic chip.

Together, these results demonstrate the possibility to operate the presented microfluidic device with low pressures (< 1 bar), as well as the relevance of using the Yeoh hyper-elastic material model to describe the deformation of the membrane. The ability to close the valves continuously could be of a great interest for many applications in order to precisely control the fluxes in the different branches of microfluidic circuits.

### 3.2 Valves validation for islets perifusion microfluidic chip

We used this model to size the valves and microchannels of a microfluidic chip dedicated to human pancreatic islets perifusion (Fig. 4a). We designed a microfluidic chip with hemicylindrical shaped microchannels of 400 μm in width and 400 μm (main channel) or 200 μm (lateral channels) in height. The aim here was to establish the orders of magnitude of pressure required to deform the membrane by the height of such microchannels. Of note, we carried out experimental tests on different valves designs (circular, square, rectangular), varying the dimensions of the pneumatic channels; we did not see any significant differences between these designs. In the presented configuration, the deformation of the membrane no longer had the shape of a symmetric spherical cap because it was laterally trapped by the fluidic microchannel walls. Nevertheless, the use of the Yeoh law was supported by the fact that a small local delamination was observed at the valves after multiple actuations at 1 bar, allowing a deformation approximated by a portion of circle of radius r (of about the height of the fluidic channel) in the longitudinal plane and of radius r/2 and in the lateral plane (Fig. 2b).

To determine the tightness thresholds of the valves experimentally, infusion of fluorescent microbeads (at Q = 10 μL/min) were recorded at different locations of the microfluidic channels (Fig. 4b). The valves were actuated at different pneumatic pressures P_p_ and the movements of the microbeads were recorded for each pressure step. The tightness of the valves was assessed at the pressure where the beads stopped moving. To do this, the consecutive frames from the raw films were subtracted using ImageJ, followed by a maximum intensity projection over 5 frames to highlight only the moving microbeads. Thus, the fluxes in microchannels were visualized by the white tracks left by the flowing microbeads, and no intensity signal were resulting from areas where the microbeads were motionless. Experiments were done with 3 valves (V_7_, V_2_ and V_3_) on 4 different microfluidic chips with similar results, and results for valves V_7_ and V_2_ are shown as examples in Fig. 4c. For the valve V7, when the valve was opened, the beads followed both path A and B. When the valve was partly opened, the beads were still flowing through path A and B, but the flow rate in path B decreased while the flow rate in path A intensified. When the valve was tightly closed, the beads flowed only through path A (Fig. 4c, top). For the valve V_2_, since there was only one possible path, the microbeads flowed along the channel for a pressure lower than the threshold pressure, and stopped flowing after the valve actuation for a pressure higher than the threshold pressure (Fig. 4c, bottom). The white arrows in the figures recapitulate qualitatively the direction of flow in each case.

Therefore, experiments showed tightness thresholds of 500 mbar for the main serpentine channel 400 μm deep (V_7_ and V_2_) and 350 mbar for the 200 μm deep lateral channels (V_3_). When applied to this configuration, the Yeoh deformation model guaranteed the tightness of the valves with working pressures above 400 mbar for the valves located in the main channel and above 250 mbar for the valves located in the lateral channels (Fig. 4d). The discrepancy between the thresholds given by the model and the experimental ones can be attributed to the fact that the complete sealing of the valves occurs at a pressure slightly higher than that the one required to deform the membrane by the exact maximum channel height. Moreover, the tightness thresholds also depend on the flow rate and the exact thickness of the membrane. For these reasons, we preferred to work with a margin ensuring the tightness of the valves under experimental conditions. These valves can be activated a large number of times without any aging being noticed. Together, these results confirmed the proper functioning of the microfluidic set-up with pressures below 1 bar.

This design resolves many conventional limitations: (1) the ability to automatically isolate and capture individual islet with the serpentine hydrodynamic trapping, (2) sufficient oxygen and dynamic glucose delivery due to continuous microfluidic perifusion, (3) detectable insulin output due to low dead volumes and flow rates. Moreover, with the hyper-elastic behavior of the membrane, the valves exhibited sufficiently large deformations to be directly integrated into the fluidic channels, without deforming the islet nor resorting to complex manufacturing methods. When opened, the valves did not induce any pressure loss in the channels. On the other hand, the valves can be closed in a perfectly tight way with pressures lower than 1 bar. Finally, using our model, the pneumatic pressure could be regulated to induce controlled flow rates in the different pathways.

### 3.3 Islets trapping and associated shear stresses

To determine the sear stress imposed to the islet during the GSIS-assay protocol, the flow in the main serpentine channel was modeled using COMSOL Multiphysics. While it has already been shown that microfluidic continuous perifusion positively affects islets function and survival**(Sankar et al., 2011)**, too high shear stress can cause cell deformation and lead to permanent damage**(Klak et al., 2021)**. To model our experimental conditions, atmospheric pressure was assigned at the inlet wall and a flow rate of 100 μL/min or 2.5 μL/min was assigned at the outlet (Fig. 5a). Indeed, the islet has to be trapped quickly at first, both to avoid sedimentation in the inlet reservoir and to allow rapid trapping. The flow rate was then reduced for the secretions phases to ensure adequate sensitivity. In both cases, the islet should not experience a too high shear stress. The valve forcing the flow toward the U-cup trap was represented by a solid block which does not allow any flow in the loop channel (path B) when activated. As described in previous studies**(Nourmohammadzadeh et al., 2013; Silva et al., 2013)**, when the valve was not actuated, the majority of the fluid flowed through path B, thus protecting the islet from shear stress (Fig. 5b). When the valve was closed (corresponding to low and high glucose stimulations), the flow was forced through path A. However, because the flow rate was significantly decreased, the associated shear stress was found one order of magnitude lower compared to the opened valve configuration (Fig. 5c). The shear values predicted on the periphery of the islet were well below the ones of > 10 Pa usually considered to cause mechanical damage**(Shintaku et al., 2008)** (Fig. 5d). A more refined analysis would have to take into account the deformability of the islet as well as its porosity, but we have sought here to establish orders of magnitude. Together, these results indicate that the islet was not subjected to problematic shear stress values during the GSIS on-chip assay. Of note, the flow rate can be continuously controlled in the loop channel (path B) thanks to the hyper-elastic membrane, so the shear stress applied to the islet can be monitored as well.

**Fig.5.**
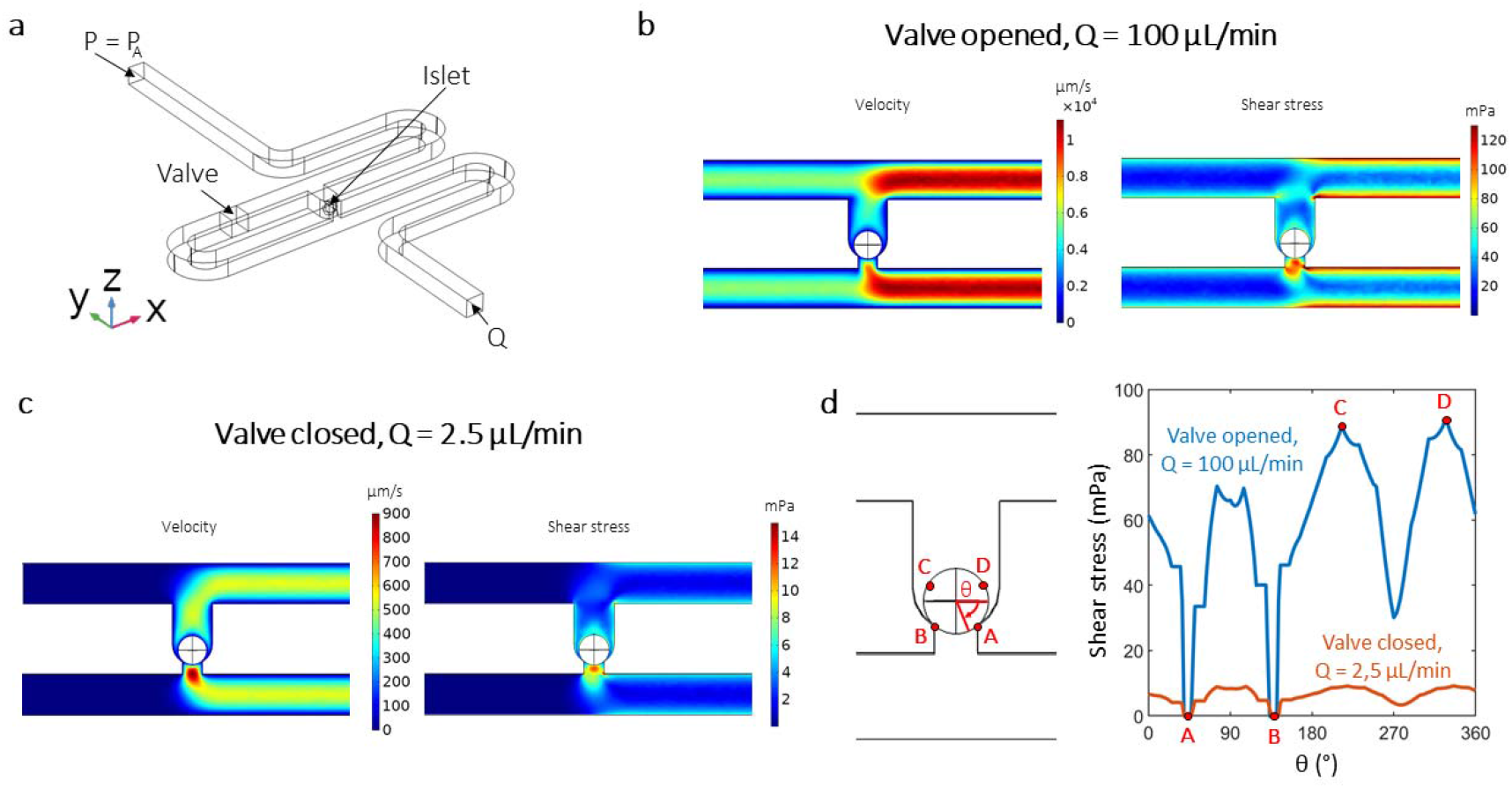
Examination of the shear stress inflicted on the islet during the GSIS assay using Comsol. a. Comsol 3D rendering of the serpentine-shaped microchannel with initial boundary conditions. b-c. Velocity and shear stress fields along the median plane of the channel near the trap site with the valve opened (b) or closed (c). d. Shear stress along the circumference line of the islet.

### 3.4 Fluidic protocol validation and islet perifusion

To validate the fluidic functioning of our microfluidic device, we performed experiments using food colorings diluted in KREBS buffer. The different possible paths for the liquid were thus visualized in white (initial filling), green (islet trapping), yellow (low glucose) and red (high glucose). In the following, for the sake of clarity, we will refer to the colored model solutions as LGS, HGS and islet retrieval respectively, in relation to the corresponding steps of the protocol in real conditions.

The experimental set-up consisted of the microfluidic chip connected to a syringepump *via* its three outlets. The chip was placed into a thermostatic chamber with a beaker of water to avoid evaporation. The valves were actuated with a pressure controller connected to the pneumatic luers ports of the chip and to a computer (Fig. 6a and Supplementary Fig. 2). The valve operation protocol was achieved using a pneumatic valve controller (LineUp P-Switch, Fluigent) connected to a pressure controller (Flow-EZ, Fluigent) and interfaced with Microfluidic Automation Tool Software (Fluigent) to fully automate the GSIS test experiments, by programming the pressurization sequence of each of the seven pneumatic microchannels during the whole experiment. Three tubings of the same length were connected to a single syringe on the syringe-pump via a T-junction tubing connector. The liquid was directed into tubing (1), (2), or (3) whether the valve V2, V3 or V6 was opened respectively (Fig 6.b). The choice of the syringe-pump was motivated by the desire to control the flow rate rather than the pressure. Indeed, during a GSIS experiment, insulin was collected in a given volume secreted by the islet during a given time interval.

**Fig.6.**
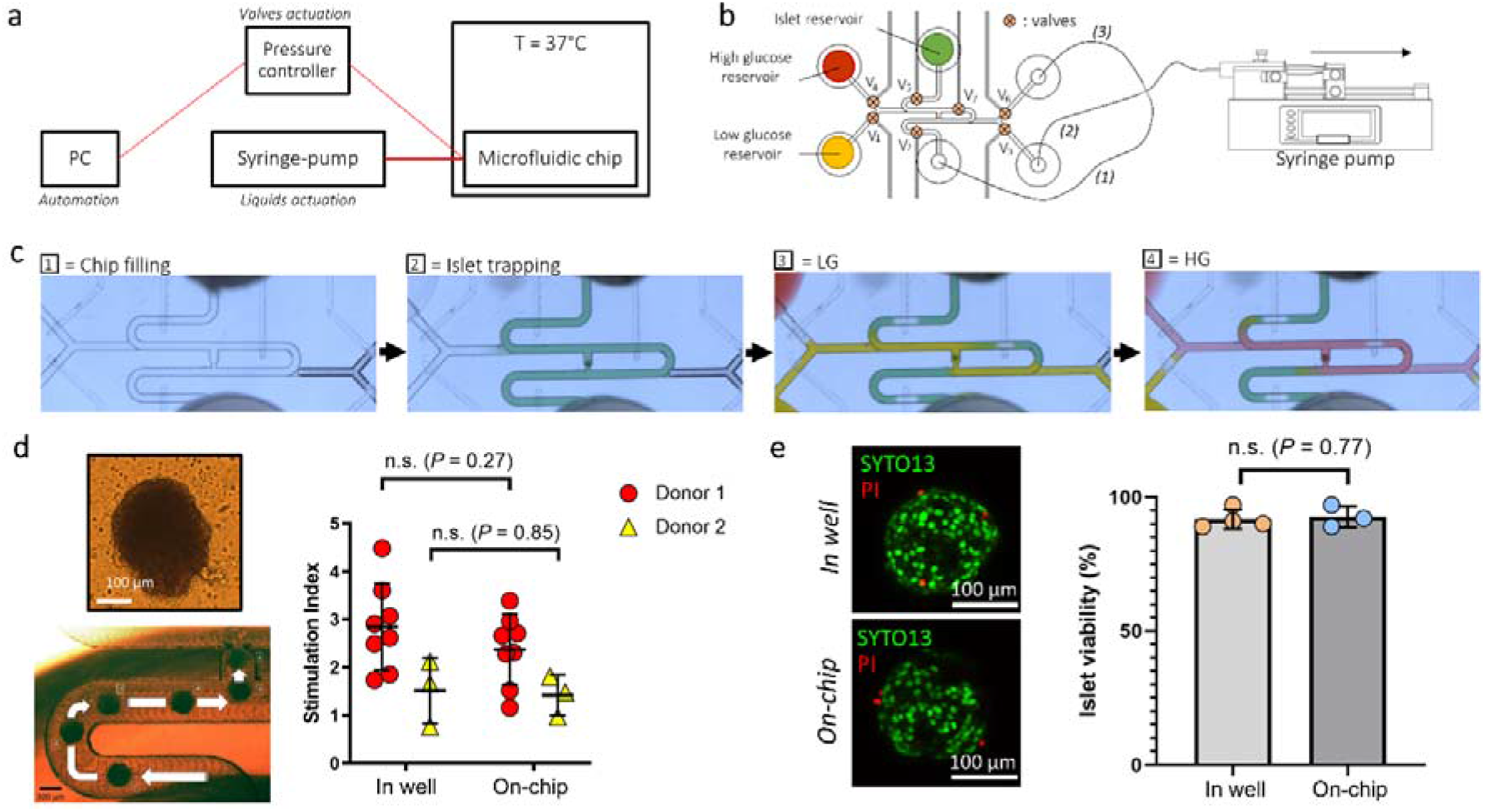
GSIS assay on-chip. a. Schematic diagram of the experimental set-up. b. Liquid actuation principle: the different reagents were introduced into the microfluidic chip using a syringe pump functioning in its withdrawal mode. The network of valves, connected to a pressure controller, allowed to direct the liquids towards different flow paths. c. Fluidic validation of the GSIS assay protocol using colored solutions: initial chip filling (1), islet loading (2), low glucose stimulation and collect (3), high glucose stimulation and collect (4). d. Representative image of the human islet obtained from cadaveric donor that is trapped (left) and comparison of the stimulation index between static (in well) and dynamic (on-chip) conditions. Each point on the graph represents the stimulation index of one pancreatic islet obtained from the measure of two samples (low and high glucose). e. Confocal z-stack maximum intensity projection renderings of a single islet stained for SYTO13 and PI after GSIS assay (left, images are representative of n=4 and n=3 islets tested in static and dynamic conditions respectively). Comparison of islet viability after GSIS assay in well or on-chip (right) Each point on the graph corresponds to one pancreatic islet. Results are shown as mean ± s.d. Unpaired t-test (two-tailed) for comparisons was conducted using GraphPad Prism 9 (GraphPad Software Inc., San Diego, CA, USA).

First, the microchannels were filled with buffer solution by capillarity. The valves V3 and V6 were closed during this process to keep the outlets clear from any liquid, all the other valves being opened (Fig. 6c.1). A human islet, obtained from a cadaveric donor, was added with a pipet into the islet loading reservoir and was flowed into the microchannel at Q = 100 μL/min through the main path (to tubing 1), the valves V1, V3, V4 and V6 being closed and V2, V5 and V7 opened. It was trapped according to the hydrodynamic principle described above. The pre-incubation phase was performed through the perfusion of LGS into the main serpentine channel using the same valve configuration *i.e*. V1, V3, V4 and V6 closed and V2, V5 and V7 opened (Fig. 6c.2). For low glucose stimulation, because once the islet was trapped, *Q_B_* > *Q_A_*, the flow was preferentially through the path B. Thus, the valve V7 was closed to force the flow through the path A to collect the secretions of the islet without being too diluted. The valves V4 and V6 were kept closed while V2 and V5 from the serpentine channel were closed. V1 and V3 were opened to achieve low glucose stimulation and supernatant collection (Fig. 6c.3). The LGS sample was collected in tubing 2. In the same way, high glucose stimulation and supernatant collection required the closing of valves V1 and V3 and the opening of V4 and V6 (Fig. 6c 4). V2, V5 and V7 were kept closed. The HGS was collected in tubing 3. At the end of the GSIS test, the islet can be retrieved for further cell culture or other assays (e.g. examination of its viability or total insulin content). To do so, the valves V1, V3, V4, V6 and V7 were closed while V2 and V5 were opened. The islet was thus pushed towards the islet reservoir using the syringe-pump in its infusion mode. All the steps of the protocol with corresponding valves openings and closings are given in Table 1.

**Table 1:** Sum-up of the valves openings and closing during the whole protocol. O = Opened, C = Closed.

| | V1 | V2 | V3 | V4 | V5 | V6 | V7 | Q<br>($\mu\text{L}/\text{min}$) |
| --- | --- | --- | --- | --- | --- | --- | --- | --- |
| Initial filling | O | O | C | O | O | C | O | / |
| Islet loading | C | O | C | C | O | C | O | 100 |
| Pre-incubation<br>(phase 1) | C | O | C | C | O | C | O | 2.5 |
| Low glucose<br>(phase 2) | O | C | O | C | C | C | C | 2.5 |
| High glucose<br>(phase 3) | C | C | C | O | C | O | C | 2.5 |
| Islet retrieval | C | O | C | C | O | C | C | 2.5 |

We proposed a biological validation of our device using human pancreatic islets. A reproducible loading of pancreatic islet into the microfluidic chip was successfully achieved *via* syringe-pump driven flow, at the flow rate of Q = 100 μL/min. As described in the sections above, an islet was precisely positioned at the trap sites without any apparent deformation, thus demonstrating a soft trapping mechanism. To validate the functionality of the islet on-chip, we adapted the conventional static culture Glucose Stimulated Insulin Secretion (GSIS) assay to the perfusion conditions of the microfluidic system by first infusing at Q = 2.5 μL/min the microchannels as described below, with 2.8 mM glucose solution for 1 h (pre-incubation phase), followed by a 1 □h of perfusion with 2.8□mM glucose solution (low glucose stimulation) and a 1 h perfusion with 16.7 mM glucose solution (high glucose stimulation). For each islet, the perfused supernatants were collected in the tubing connected to the exit ports for low and high glucose conditions, resulting in two 150 μL samples (low and high glucose) for further ELISA analysis. The insulin secreted by the islet on-chip increased significantly from the first incubation of the low glucose condition to the high glucose condition. Pancreatic islets obtained from InSphero (donor 1) showed a stimulation index (SI) of 2.8 ± 0.9 and 2.4 ± 0.7 in static and on-chip conditions respectively (mean ± sd, n=8 for each condition). Pancreatic islets obtained from Montpellier Hospital (donor 2) showed a SI of 1.5 ± 0.7 and 1.4 ± 0.4 in static and on-chip conditions respectively (mean ± sd, n=3 for each condition). Overall, the SI results were similar between both the static cultures and the on-chip experiments, thus validating that glucose response from the islets was not impaired in our microfluidic device (Fig. 6d). Moreover, SI above 1 demonstrate the correct functionality of islets, and therefore that the Ecoflex does not induce toxicity. Although this last point has already been proven in various studies**(Luis et al., 2019; Salvatore et al., 2017; Wang et al., 2012)**, we performed viability tests to ensure the non-toxicity of our device and protocol on the pancreatic islets. Viability tests were conducted after GSIS assay in well (n=4 individual islets, donor 1) and on-chip (n=3 individual islets, donor 1) *via* SYTO13 and Propidium Iodide (PI) double staining (Fig. 6e). Viability remained above 90% after GSIS assay in both static and on-chip conditions, with no significant difference found between the two conditions. This also proves the biocompatibility of our device.

All together, these results demonstrate the ability of the presented microfluidic device to direct fluxes easily using a network of valves that can be integrated in the fluidic circuit. The device we developed has the advantage of enabling to perform a GSIS assay on one single islet on-chip with similar results to the classical static conditions. The islet trapping function in the device allows an automatic positioning of the islet, but also, by reversing the direction of the flow, allows to retrieve the islet for offline analysis after on-chip experiments. Compared to conventional methods where numerous manual operations are needed, our microfluidic platform allows to perform a GSIS assay in a fully automated way without requiring complex technical settings. In addition, because a microfluidic approach generates very low dead volumes, insulin secretion kinetics of individual islet could be determined by collecting samples at short time intervals. Regenerative medicine with islet transplantation, a cell therapy for diabetes, has emerged for a decade as a promising effective treatment to restore good glycaemic control in insulin independent diabetic patients with severe form of diabetes**(Lablanche et al., 2018)**. The device we present here could help to quickly test and sort the best islets before transplantation. It has already been shown that this serpentine architecture can be parallelized to trap tens of islets**(Zbinden et al., 2020)**, thus our device could be easily adapted for higher throughput for applications requiring it. Finally, this microfluidic platform can be used for diverse biological assays and should enable the analysis of diverse secretomes. While the presented device does not embed insulin sensing on-chip yet, this hyper-elastic membrane based technology has already been proven reliable for quantitative biological assays in one of our previous works**(Parent et al., 2018)**. Some investigations are ongoing in our labs to address this and to develop a fully automated and integrated microfluidic platform for the assessment of pancreatic islets function.

## 4. Conclusions

In this paper, we present a new microfluidic platform that provides straightforward solutions to overcome the limitations often encountered with conventional pancreatic islets perifusion systems. Managing to decrease the number of pancreatic islets down to one islet and the ease of use of our device is of great interest for clinical use. Indeed, our device could enable an extra delivery criteria for grafting pancreatic islets to patients. The trapping of individual islets as well as the temporal and spatial control of the fluxes were automated through a network of hyper-elastic valves easy to integrate on-chip. Moreover, the chip was manufactured in COC for scalable manufacturing purposes. By altering the number or the arrangement of lateral microchannels, one can customize the device for a wide range of biological assays. The microfluidic chip can be improved by integrating in series either the U-cup shaped trap to experiment on a pool of several islets, or the whole unit presented here to study individual islets at a higher throughput. Of note, parallelizing the traps while varying their dimensions can be of interest to sort islets by size. Also, integrating a readout on-chip such as an ELISA for insulin sensing**(Glieberman et al., 2019)** or other micro-sensors**(Chmayssem et al., 2021)** would be particularly beneficial. Moreover, to reduce the footprint of the overall device, the syringe pump could be replaced by a peristaltic pump on-chip functioning on the principle of the stretchable membrane. Finally, this work is part of a more general framework in which microphysiological systems can incorporate a higher physiological relevance compared to static cultures, such as a functional vasculature**(Rambøl et al., 2019)**. These model systems will benefit from being handled with microfluidic techniques, which allow a more direct access to the biological tissues and a precise control of the stimuli.

## Author Contributions

Conceptualization: C. Q., Y. F., F. N., X. G., methodology: C. Q., Y. F., X. G., project administration: F.N, X.G., validation: C.Q., E.T., J-L. A., Y.F., X.G., investigation: C. Q., E. T. formal analysis: C. Q., E. T., writing original draft: C.Q., writing review & editing: Y. F., E.T., J-L. A., F.N, X. G. Supervision: Y. F., J-L. A., F.N, X. G., funding acquisition: F. N., Y. F., X. G.

## Acknowledgements

We thank F.Boizot and M.Alessio for the manufacturing of microfluidic chips. We thank the French Carnot Institute and CEA for funding.

