## Supplementary Material for "Microfluidic device integrating a network of hyper-elastic valves for automated glucose stimulation and insulin secretion collection from a single pancreatic islet"

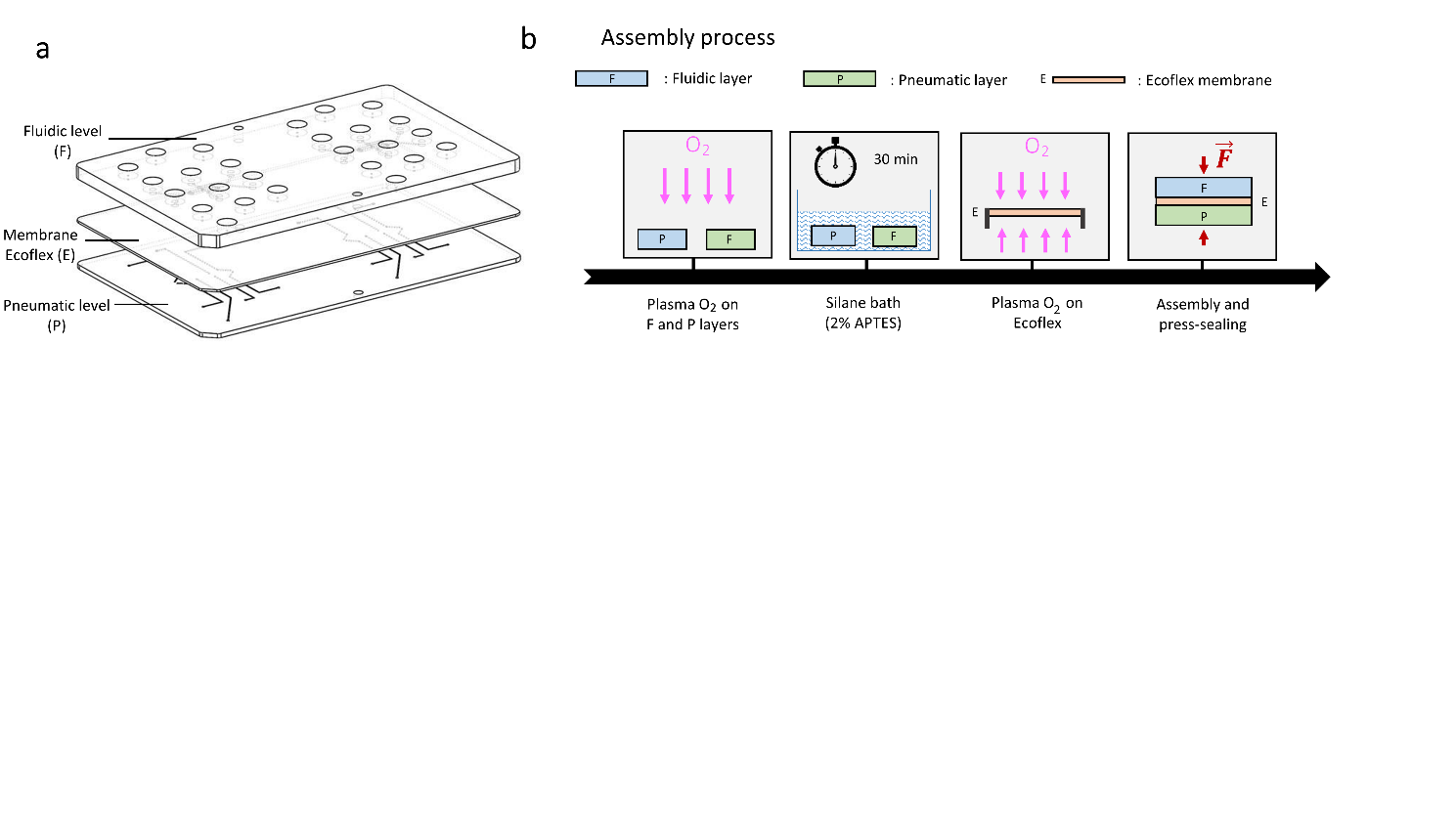

Supplementary Fig. 1. Assembly of the microfluidic chip. a. Solidworks 3D exploded view of the GSIS assay microfluidic chip showing the three layers to assemble: the fluidic level (F), the pneumatic level (P) and the Ecoflex stretchable membrane (E) sandwiched between the two. b. Chip assembly process, step by step (see Material & Methods for details).

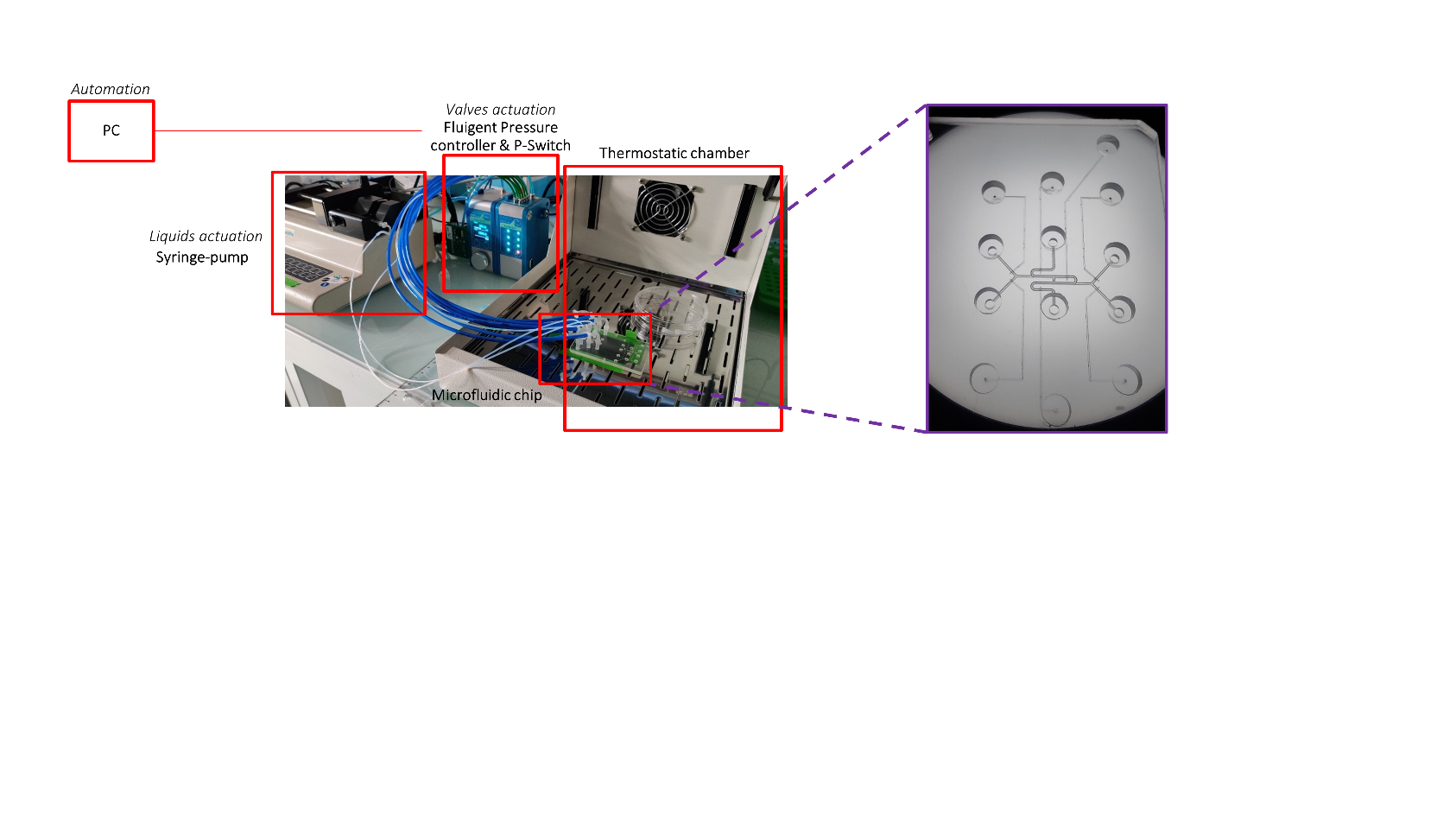

Supplementary Fig. 2. Experimental set-up for automated GSIS assay on-chip. Photograph of the experimental set-up showing the different elements of the microfluidic platform, with a detailed view of the microfluidic chip (inset).

Supplementary Note 1: Trapping principle and islets loading

The microfluidic channel was composed of a serpentine-shaped loop channel with a narrow U-cup-shaped region that functioned as a 3D-cell aggregate trap (Fig. 1b). Using the Darcy-Weisbach equation, we obtain the pressure difference $\Delta P=f_{D}\frac{L}{D} \rho\frac{v^{2}}{2}$ (where the hydraulic diameter is $D=\frac{4A}{P}$ with $A$ the cross-sectional area of the flow and $P$ the wetted perimeter of the cross-section, $L$ is the length of the considered channel, $f_{D}$ is the Darcy friction factor, *ρ* is the density of the fluid, $v$ the mean flow velocity). The Darcy friction factor is related to the Reynolds number $Re =\frac{\rho vD}{\mu}$ (where $\mu$ is the dynamic viscosity of the fluid) and the aspect ratio $\alpha$ (0$\leq\alpha\leq1$) by $f_{D}Re=C\left( \alpha\right)$ where denotes a constant that is a function of the aspect ratio. It follows $\Delta P=\frac{C\left( \alpha\right)}{32} \frac{{\mu L Q P}^{2}}{A^{3}}$ where we introduced the flow rate $Q = vA$. Thus, the ratio of the flow rates along the U-cup path (A) and the loop path (B) is given by:

$$\frac{Q_{A}}{Q_{B}}=\left( \frac{C_{B}\left( \alpha_{B} \right)}{C_{A}\left( \alpha_{A} \right)} \right)\left( \frac{L_{B}}{L_{A}} \right)\left( \frac{P_{B}}{P_{A}} \right)^{2}\left( \frac{A_{A}}{A_{B}} \right)^{3}$$

In the case of rectangular sections, $A = W H$and $P = 2 (W+H)$, and the design should be such that an islet flowing through the serpentine channel is delivered to the trapping region where the hydrodynamic resistance is initially lower compared to the loop path. The operating criterion of the trap is therefore as follows:

$$\frac{Q_{A}}{Q_{B}}=\left( \frac{C_{B}\left( \alpha_{B} \right)}{C_{A}\left( \alpha_{A} \right)} \right)\left( \frac{L_{B}}{L_{A}} \right)\left( \frac{W_{B}+H}{W_{A}+H} \right)^{2}\left( \frac{W_{A}}{W_{B}} \right)^{3}>1$$

As a result, islets in the flow were carried into the U-cup shaped constriction. The loading of different components was done through a syringe pump connected to the outlet of the channel in its withdrawal mode. Islets stimulation and secretions collection were achieved through the lateral channels housing at the entrance two reservoirs for low and high glucose solutions, and connected to the syringe-pump at the outlets.

Supplementary Note 2: Modelling of the hyper-elastic membrane deformation

We modelled the deformation of the hyper-elastic membrane using the Yeoh law, a higher-order extension of the neo-Hookean law:

$$\sigma_{Y}=2\left( \lambda^{2}-\frac{1}{\lambda^{4}} \right)\left( c_{1}+2 c_{2}\left( 2\lambda^{2}+\frac{1}{\lambda^{4}}-3 \right)+3 c_{3}{\left( 2\lambda^{2}+\frac{1}{\lambda^{4}}-3 \right)^{2}}^{2} \right)$$

In these relations, the $C_{i}$ are constants specific to each material. It can be noted that, in general, we distinguish elongations $\lambda_{1}$, $\lambda_{2}$ and $\lambda_{3}$ in the main directions of space. In the equibiaxial case studied here, $\lambda_{1}=\lambda_{2}=\lambda$ . Finally, to this equation must be added an initial stress on the membrane that exists at the time of assembly of the microfluidic chip. Moreover, the formula of the Laplace pressure, applied by analogy to the membrane thickness is $\Delta P=\frac{2e}{R}\sigma$ where the stress $\sigma$ plays the role of the surface tension. The geometry of the device shown in the inset of Fig. 2b led to the relation $R=\frac{r^{2}+h^{2}}{2h}$. Moreover, since the Ecoflex is incompressible, we can write the incompressibility hypothesis $e=\frac{e_{0}}{\lambda^{2}}$. We thus obtain $\Delta P=\frac{4e_{0}h}{r^{2}+h^{2}}\frac{1}{\lambda^{2}}\sigma$ with $\lambda=\frac{R\theta}{2r}=\frac{r^{2}+h^{2}}{2rh}\sin^{-1} (\frac{2rh}{r^{2}+h^{2}})$. The volume of the chamber (spherical cap) being related to $h$ and $r$ by $V=\frac{\pi h^{2}\left( 3r-h \right)}{3}$, this gave us the relation between the volume $V$ of liquid introduced and the pressure difference $\Delta P$ between both sides of the membrane, directly related to the deformation height of the membrane thus to the flow passage section.
